# No evidence of sexually antagonistic coevolution in Drosophila reproductive tract transcriptomes

**DOI:** 10.1101/2025.05.07.652506

**Authors:** Rachel C. Thayer, Elizabeth S. Polston, David J. Begun

## Abstract

Drosophila seminal fluid proteins (SFPs) are often cited as an example of interlocus sexual conflict, wherein the proteins increase male fitness while decreasing female fitness, spurring recurring female counter adaptations and rapid molecular evolution. This model predicts that male-expressed genetic variation in the accessory gland, which produces seminal fluid, should generate counter-evolving genetic pathways in females, resulting in sexual coevolution. Using a trio of *D. melanogaster* populations exhibiting substantial SFP expression divergence due to recent selection, we test for coevolution in the female post-mating transcriptome in the lower reproductive tract and head. Contrasting predictions of sexual antagonism, female postmating gene expression is indifferent to male population of origin. Instead, our results better support the alternative hypotheses that environmental variation is the source of selection on male SFP gene expression, and that population differentiation in the female post-mating transcriptome is generated by female-expressed genotypic differentiation.

## INTRODUCTION

A central goal of evolutionary genetics is to identify the selective forces responsible for patterns of adaptive molecular evolution. Drosophila Seminal Fluid Proteins (SFPs, (Wigby et al. 2020) are often presented as a leading case study for postcopulatory, intersexual conflicts driving rapid gene expression and protein sequence evolution. This model posits that after their transfer to the female reproductive tract (FRT), SFPs manipulate female post-mating physiology to advance male fitness, such as by delaying remating, inducing over-investment in egg-laying, or abbreviating lifespan (Holland and Rice 1998; Rice 1996; Hollis et al. 2019; Chapman et al. 1995; Lung et al. 2002), reviewed in (Hopkins and Perry 2022). Females would counter-evolve resistance mechanisms, generating sustained selection on the associated genes and proteins. This dynamic is one proposed cause for the rapid SFP evolution observed across many taxa (Begun and Lindfors 2005; Wyckoff, Wang, and Wu 2000; Swanson, Nielsen, and Yang 2003; Swanson and Vacquier 2002).

Previous work connects SFP dosage to female post-mating physiology and gene expression, but how these genetic interactions may shape sexual coevolution is disputed (Hopkins and Perry 2022). In terms of physiology, experimentally-generated hypomorphic and null SFP alleles have been used to reveal that certain SFPs have a variety of effects on post-mating outcomes (Chapman and Davies 2004; Chapman et al. 2003; Wolfner 2002), including effects on female gene expression (McGraw, Clark, and Wolfner 2008; Gioti et al. 2012). Experimental evolution work that eliminated female coevolution (Rice 1996) or sexual selection (Hollis et al. 2019) encouraged the interpretation that these SFP-female interactions could translate into a broader antagonistic evolutionary dynamic, based upon the concurrence in these experiments of male fitness improvements via sperm competition advantages, possible female fitness reductions via shortened lifespans, and differences across reproductive transcriptomes in both sexes (Hollis, Houle, and Kawecki 2016; Hollis et al. 2014). Considering natural evolution among Drosophilid flies, gene expression in the female reproductive tract is perturbed by mating to a heterospecific versus conspecific partner (Ahmed-Braimah, Wolfner, and Clark 2021; Bono et al. 2011), which may factor into reproductive isolation, and must begin as within-species population differentiation in male-female genetic interactions (Lollar et al. 2023; Alipaz, Wu, and Karr 2001; Garlovsky et al. 2020), such as SFP-female interactions (Wensing and Fricke 2018). Attempts to associate natural genotypic variation in male *melanogaster* with whole-female post-mating transcriptome variation have yielded mixed findings (McGraw et al. 2009; Delbare et al. 2017), leaving both the potential coevolutionary interactions and the identities of prospective female-expressed, counter-evolving loci still unclear. An approach using strategically selected male genotypes associated with both SFP dosage variation and adaptive evolutionary context, and focusing more sensitively on select female organ transcriptomes, could be informative.

Here, we explicitly test for coevolution of reproductive tract gene expression using natural *D. melanogaster* populations from the well-studied North American latitudinal cline (Adrion, Hahn, and Cooper 2015) and a population sample from the species’ ancestral range in Zambia (Pool et al. 2012). In the male accessory gland, the primary source of seminal fluid, Panama males exhibit highly diverged gene expression from males from Maine, with a significant enrichment for differentially expressed SFP genes (Fig. 1A) (Cridland, Contino, and Begun 2023). Most of this divergence appears to be the result of evolution in the Panama population. The SFP gene expression differentiation is starkly asymmetric, with Panama males exhibiting naturally hypomorphic expression of 48 SFPs relative to Maine males (∼16% of *D. melanogaster* SFPs), strongly suggestive of a role for selection in driving the expression divergence. More generally, genetic differentiation (Fst) along the N. American east coast cline is low (Turner et al. 2008; Reinhardt et al. 2014), supporting the interpretation that most latitudinal phenotypic differentiation results from recent spatially varying selection. Transcriptome comparisons of North American vs. Zambia populations revealed >2300 differentially expressed accessory gland genes, including 104 SFP genes (27% of accessory gland genes and 35% of SFPs differentially expressed, Fig. 1B). We hypothesized that under the sexual conflict model this recent, adaptive male differentiation would correspond to diverged male-female interactions. For example, Zambia females mated to Panama males might enact a more Panama-like transcriptional response, or more generally an aberrant response, consistent with the idea that recent selection on accessory gland transcriptomes has changed the composition of seminal fluid and its influences on female transcriptomes.

**Fig 1:**
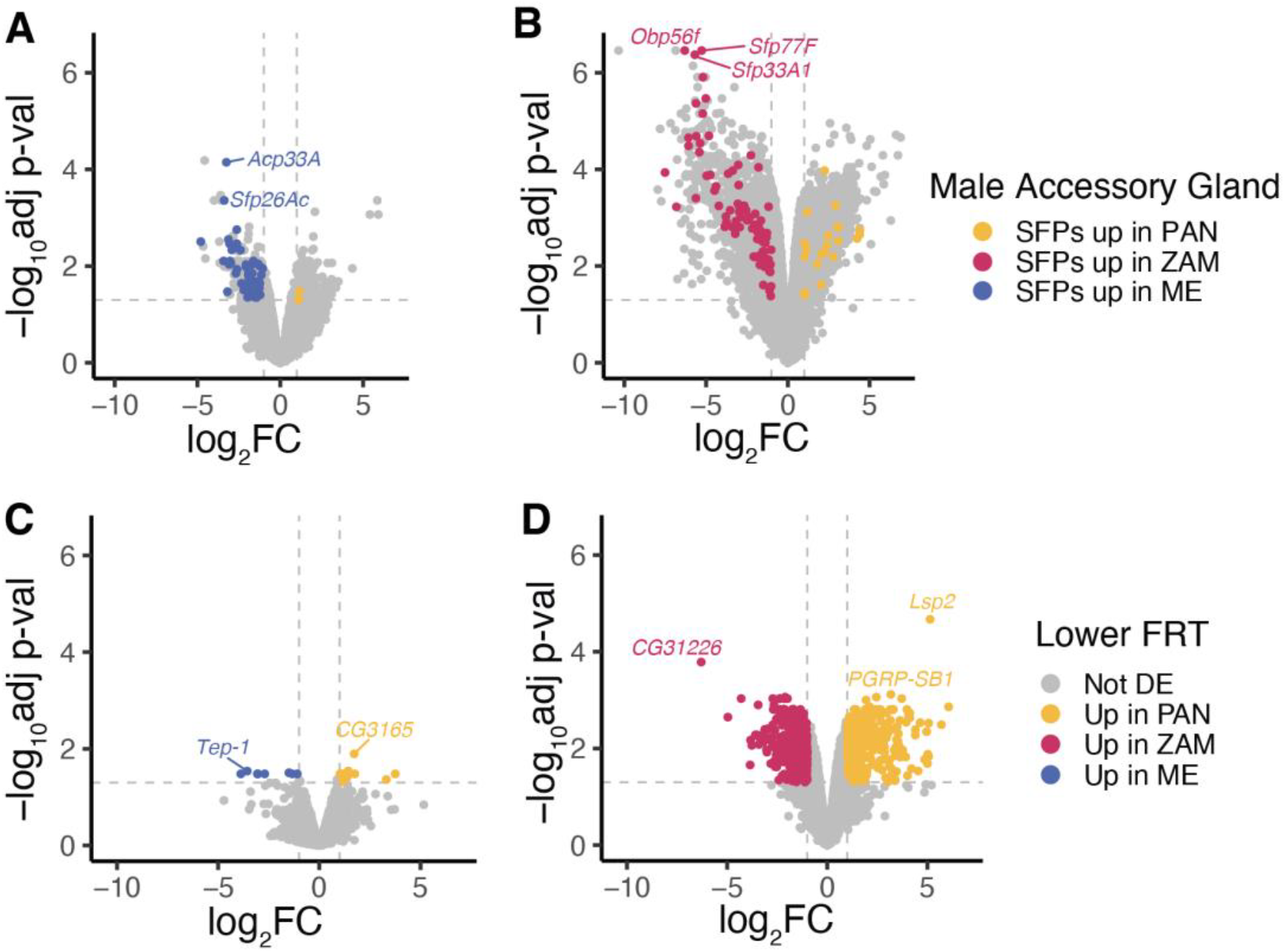
Reproductive transcriptome population differentiation between Panama, Maine, and Zambia. *A-B)* Male accessory gland transcriptomes; differentially expressed SFPs are colored according to the higher-expressing population: red for Zambia, yellow for Panama, blue for Maine. *A)* Maine versus Panama accessory gland. *B)* Zambia versus Panama accessory gland. *C-D)* Mated-state lower reproductive tract transcriptomes in females mated to a same-population mate. Differentially expressed genes are colored according to the higher-expressing population: red for Zambia, yellow for Panama, blue for Maine. *C)* Maine versus Panama FRT. *D)* Zambia versus Panama FRT.

To test for male effects on female transcriptome variation, we measured female post-mating gene expression following mating to either a co-evolved same-population male, or a diverged male (Fig. 2). We measured gene expression in the somatic female reproductive organs that directly interact with semen (i.e. the uterus, sperm storage organs, reproductive glands, and common oviduct). Gene expression in these organs is strongly mating responsive over a well-defined time course (Mack et al. 2006; McDonough-Goldstein et al. 2021), providing a high dimensional assay to quantify the effects of adaptive male transcriptome evolution on females and simultaneously identify interacting female genetic pathways. Beyond the effects of SFP expression variation, these experiments also capture any other effects of population divergence on the composition of seminal fluid, such as divergence of SFP sequences, and any possible variation in the concentrations of amines, lipids, and sugars in seminal fluid.

**Fig 2:**
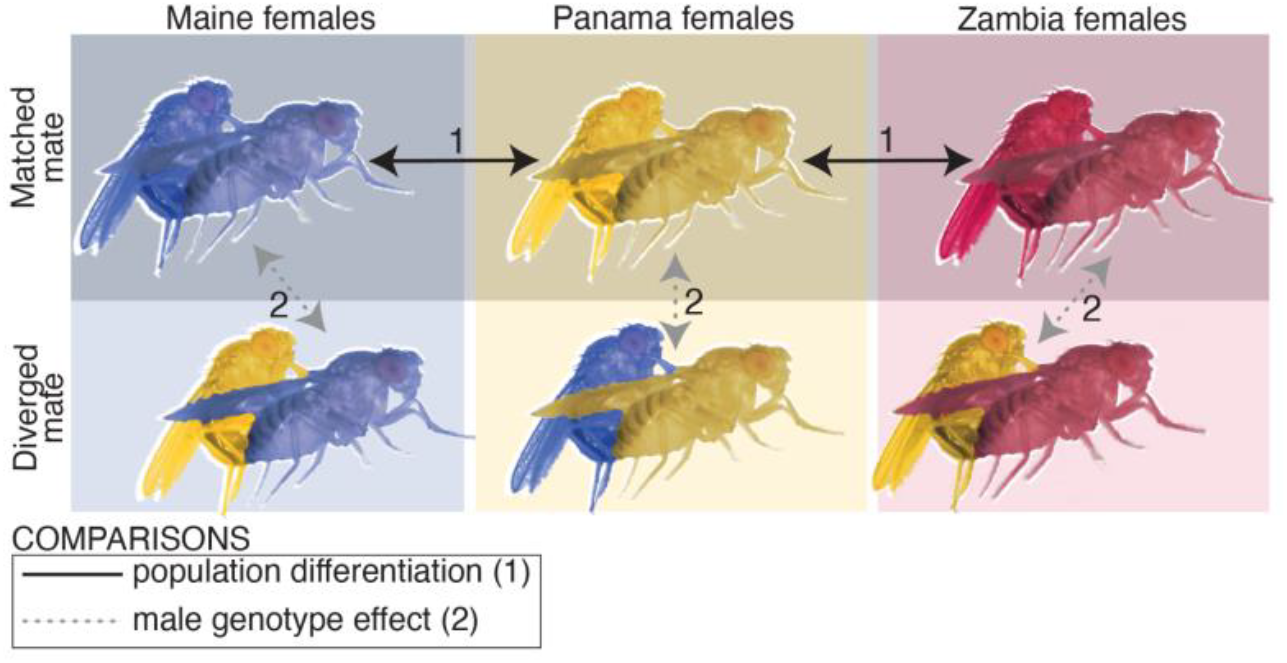
Experimental strategy using crosses among three populations to distinguish inter-population differentiation and male genotype effects.

## RESULTS

Female reproductive tract transcriptomes were differentiated among populations proportionally to population relatedness, with greater differentiation between Zambia versus the American populations (Fig. 1 C-D). Regarding co-evolutionary dynamics, the most important result is that male geographic origin had little detectable effect on FRT transcriptomes (Fig. 3). Among >8100 FRT-expressed genes, only three genes were differentially expressed between females mated with foreign vs. same-population males (Fig. 3 A-C) which is no more than expected by chance given multiple hypothesis testing. Consistent with the conservative interpretation that they are false positives, these three genes exhibited modest log fold changes. Comparing population differences for FRT gene expression of females mated with males from their own population, we identified 21 differentially expressed genes between Maine and Panama (Fig. 1C) and 884 differentially expressed between Panama and Zambia (Fig. 1D). Together, these findings indicate that female genotype is substantially more important to interpopulation differences in the mated-state FRT transcriptome than either male genotype or male-female genotypic interactions, which appear negligible.

**Fig 3:**
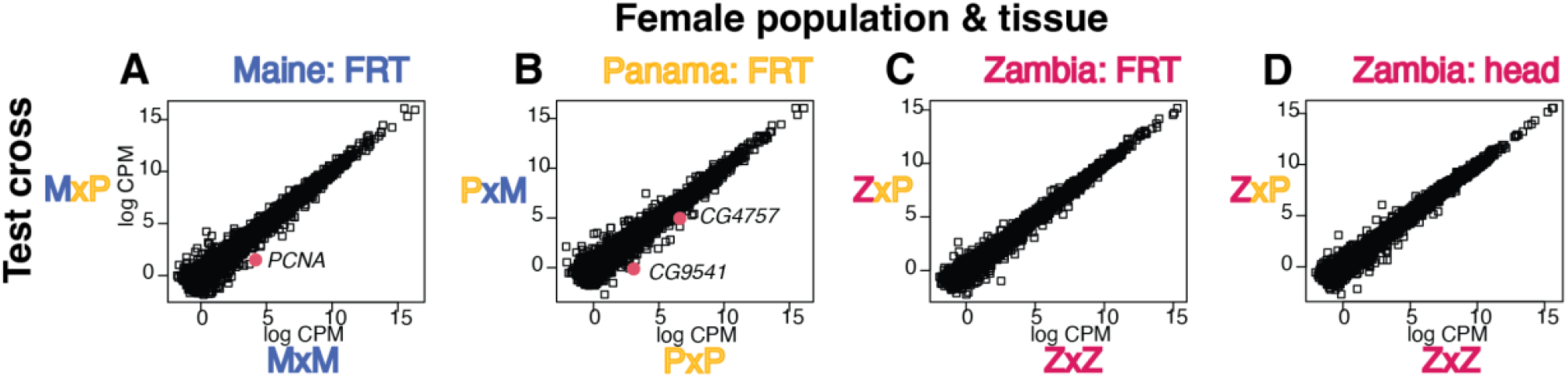
Transcriptomes from inter-population test crosses. *A-D)* Mean log Counts Per Million in females with a matched-population mate (x-axis) versus a diverged mate (y-axis). Data points for significantly differentially expressed genes are large red points with the gene name labeled. *A)* FRT in Maine females with a Maine versus a Panama mate. *B)* FRT in Panama females with a Panama versus Maine mate. *C)* FRT in Zambia females with a Zambian versus Panama mate. *D)* Head of Zambia females with a Zambian versus Panama mate.

Because reception of certain SFPs induces various female behaviors, (Isaac et al. 2010; Chen et al. 1988; Bath et al. 2017; Scheunemann et al. 2019), and because a prior study comparing whole-fly, mated-state female transcriptome divergence highlighted some female-head-expressed genes (Delbare et al. 2017), we also screened for effects of male population divergence on gene expression in the female head. Using the most diverged pairwise comparison for the accessory gland transcriptome, Panama and Zambia (Cridland, Contino, and Begun 2023), we recovered no male population effect on the female head transcriptome (Fig. 3D). Thus, across a set of organs with hundreds of mating-responsive genes (Dalton et al. 2010; McDonough-Goldstein et al. 2021) – including the tissues that directly contact sperm and semen, and that mediate wide-ranging behavioral post-mating responses – females exhibit effectively identical post-mating transcriptomes regardless of substantial accessory gland divergence of their mates.

## DISCUSSION

Using populations with evidence of adaptive divergence in male accessory gland and SFP gene expression, we find no evidence that this divergence affects the reproductive tract or head transcriptomes of their mates. While it remains possible that accessory gland adaptations have some manipulative influence that leaves no signature on female transcriptomes, our results suggest it is unlikely that males regulate many granular aspects of female physiological variation via many individual components of semen. The fact that diverse aspects of female post-mating biology are altered by male knockdown of certain individual SFPs may indicate only that these specific SFPs–primarily Sex Peptide–are cues *melanogaster* females use to recognize that mating has occurred. Upon receiving these minimal cues, females hang their own regulatory scaffolding for enacting their post-mating physiology, including its transcriptomic components. Females may often use minimal cues to initiate reproductive state transitions. For example, in mice, mechanical stimulation is sufficient to induce pseudopregnancy (Stone and Srodulski 2023), which can similarly be described as a collection of many physiological and behavioral changes; this does not indicate that male mice use physical stimulation to regulate each discrete component of pseudopregnancy. Similarly in flies, it may be mischaracterizing the power dynamic–and by extension, the evolutionary implications– to describe Sex Peptide and other SFPs as vehicles by which males regulate female immunity, behavior, gene expression, and metabolism.

If not to exert control over female physiology, what selective forces may drive adaptive inter-population SFP expression differentiation (Cridland, Contino, and Begun 2023)? Latitudinally varying environmental factors that may influence sperm viability, storage or competition are obvious possibilities. SFPs are components of fluid that is, effectively, a cell culture media for spermatozoa, both in seminal fluid and in the female storage organs (Thayer et al. 2024); perhaps the ‘recipe’ needs optimization across different temperatures (van Heerwaarden and Sgrò 2021; Gandara and Drummond-Barbosa 2023; Scossiroli 1954). Variation in seasonality, UV exposure (Svetec et al. 2016), and ecological community composition may also be relevant. Sperm competition among males is another prospective source of selection on SFP sequence and expression (Fiumera, Dumont, and Clark 2005, 2007; Patlar and Civetta 2022). Some experiments associate sperm competition advantages with collateral damage to females (Rice 1996; Hollis et al. 2019; Civetta and Clark 2000), implying that sperm competition adaptations are inherently also sexually antagonistic, and should still generate female counter-adaptations. Because we find no evidence of antagonistic coevolution in reproductive tract transcriptomes, our data are more easily compatible with a model wherein any possible sperm competition adaptations among these populations are not strongly deleterious to female fitness (Jiang et al. 2011; Castrezana et al. 2017), either because the laboratory-based findings do not reflect phenomena in natural populations, or because female fitness and lifespan not strongly correlated.

## MATERIALS AND METHODS

### Fly samples

Flies were sampled from 30 isofemale lines that were established from inseminated females collected in Fairfield, Maine (September 2011) and Panama City, Panama (January 2012) (Zhao et al. 2015). We also sampled flies from 10 isofemale lines from Zambia (Pool et al. 2012). 3-7 day old, virgin females were introduced to three males in a vial and watched to confirm copulation. Males of mixed ages were held in vials in same-genotype groups of five for five days prior to the experiment, ensuring that none had recently mated. After mating, males were removed and females were rested for 4-6 hours before dissection, with the same range of rest times evenly represented among replicates and treatments. Flies were housed on standard yeast-cornmeal-agar food at 25° C on a 12-hour light:dark cycle. We sampled 15 flies from different isofemale lines and pooled for each North American population (i.e. 15 flies per population sample, with 15 genotypes equally represented). For Zambia, we used 1 fly from each of 10 isofemale lines per sample. We used three replicates per treatment, however, because Zambia line 110 females would not readily mate with Panama males, the Zambia x Panama treatments had only 9 flies in each replicate.

### RNA isolation and sequencing

The lower FRT of mated females was dissected from the common oviduct to the posterior uterus, inclusive of the sperm storage organs, female accessory glands, and reproductive-associated fat body. RNA extraction and sequencing methods followed (Cridland, Contino, and Begun 2023). Paired-end, 150-base pair reads were generated on an Illumina NovaSeq 6000. One Zambia x Zambia head replicate failed RNA quality screening and was dropped from further analysis.

### Analysis

Differential expression analysis followed (Cridland, Contino, and Begun 2023). Reads were aligned to genome v6.41 (downloaded 2021 August 9 from Flybase, (Jenkins et al. 2022) with hisat2 v2.1 (Kim, Langmead, and Salzberg 2015). Read counts were generated using featureCounts v2.0.3 (Liao, Smyth, and Shi 2014) and differential expression was called using limma 3.50.3 (Ritchie et al. 2015) within R 4.1.2. To be considered expressed, genes required a median TPM ≥ 1 in at least one treatment per pairwise comparison. To be called differentially expressed, we required a logFC with absolute value >1 and adjusted p value < 0.05.

## Data availability

Sequences are hosted at NCBI SRA under PRJNA1257015.

## ACKNOWLEDGEMENTS

We thank Giovanni Hanna for contributing to fly dissection and Julie Cridland for helpful advice. This work was supported by the National Institute of General Medical Sciences via R35GM134930, R35GM156525, and F32GM146419, and by the National Institute of Child Health and Human Development via K99HD115833.

## Notes

### Competing Interest Statement

The authors have declared no competing interest.

